# One Health assessment of antimicrobial-resistant Enterobacterales and ESKAPE pathogens in little stints (*Calidris minuta*) and aquatic ecosystems of the Kenyan Rift Valley

**DOI:** 10.1101/2025.10.26.684674

**Authors:** Catherine W. Mbuthia, Rael J. Too, Alexander Mzula, Titus S. Imboma, John Kiiru, Samuel Kariuki, Abubakar S. Hoza

## Abstract

Palearctic migratory little stints (*Calidris minuta*) can acquire resistant bacteria from anthropized environments and spread them across different hosts and borders. This two-year cross-sectional study assessed the prevalence of multidrug-resistant (MDR) and extended-spectrum beta-lactamase (ESBL)-producing Enterobacterales and ESKAPE pathogens isolated from *C. minuta* and their aquatic foraging ecosystems at two Kenyan Rift Valley lakes, Bogoria (low anthropogenic activities) and Magadi (high impact). A total of 184 fecal samples from *C. minuta* and 48 water samples were collected during the birds’ arrivals (cohort 1) and departures (cohort 2). Samples were cultured, bacterial isolates were identified using MALDI-TOF MS platform and tested against 12 antimicrobials using the Kirby-Bauer disk method. Of the 294 isolates (16 genera and 33 species), Enterobacter species (31%) and *Escherichia coli* (17.3%) were predominant. Resistance was highest for ampicillin (50%), amoxicillin-clavulanic acid (36.4%), and tetracycline (32.7%), and lowest for meropenem (1.0%) and cefepime (3.4%). The predominant MDR phenotype was a combination of resistances to ampicillin, tetracycline, and trimethoprim-sulfamethoxazole. Enterobacter species showed the highest frequencies of MDR (8.5%) and ESBL-MDR (4.4%) phenotypes, while Acinetobacter species were the most frequent ESBL producers. Despite observing higher median resistances in isolates from Lake Magadi (7.1), *C. minuta* (8.0) and cohort 2 (7.1) than those from Lake Bogoria (6.7), water samples (5.9) and cohort 1 (6.5), these differences were statistically insignificant (p-values= 0.833, 0.147, 0.210). This suggests that while human activities drive AMR spread, resistant strains are pervasive even in minimal human-influenced environments. This is the first study to link *C. minuta* to the AMR epidemiological circuit. Our findings underscore the need to include migratory wild birds in AMR surveillance in the Kenyan Rift Valley, to implement stringent environmental stewardship measures to curb anthroponotic AMR transmissions and utilizing whole-genome sequencing to accurately trace the origin and dissemination pathways of AMR strains.

## Introduction

Antimicrobial resistance (AMR) poses a critical and multi-faceted threat to global health, with impacts spanning human, livestock, wildlife, and environmental domains (Ahmed et al., 2024). The pervasive detection of AMR at the interfaces between humans, animals, and the environment underscores the necessity of the One Health framework for mounting an effective response (2) (3). Although the natural environment represents AMR’s primordial reservoir (4), anthropogenic activities are the primary drivers of its current acceleration (5). Key among these are the intensive use of antimicrobials in agriculture (6) (7) (8), industrial pollution (9), and their misuse in clinical (10) and veterinary settings (11). These human-derived pressures inevitably impact wildlife, with migratory birds demonstrating high frequencies of AMR strains linked to contaminated habitats (12) (8) (13).

Within the One Health context, migratory birds are increasingly recognized as potential global vectors and sentinels for AMR (14). Their long-distance movements across continents (15) (16) (13) bring them into contact with a spectrum of anthropogenically influenced landscapes, from farmlands (8), water bodies (17) (18), to landfills (19) and urban centers (20), which serve as hotspots for the exchange of resistant bacteria.

The little stint (*Calidris minuta*) exemplifies this phenomenon. This small shorebird, with a population of approximately 1.6 million (21), undertakes a vast annual migration from its Arctic breeding grounds to wintering sites across Africa and Asia (22) (21). Its reliance on wetland ecosystems, such as the Rift Valley lakes of Kenya, and its extended residency in these areas (23), create significant potential for the acquisition and dissemination of resistant bacteria.

The public health significance of this dynamic is heightened by the specific bacteria being isolated. The detection of resistant Enterobacterales and notorious ESKAPE pathogens in wild birds (24) (25) (26) is particularly alarming, as these groups are frequently associated with multi-drug resistance (MDR) in clinical settings. Resistance to last-line agents like third to fifth generation cephalosporins and carbapenems is often mediated by genes for extended-spectrum beta-lactamases (ESBLs) and carbapenemases. Critically, these genes are commonly carried on mobile genetic elements, facilitating their dissemination across bacterial species and thereby intensifying the global AMR crisis (27).

While anthropogenic activities are recognized as a key driver of AMR dissemination by migratory birds globally, data from Kenya on this phenomenon remain scarce. This study aimed to address this critical gap by evaluating the influence of human activity on AMR prevalence in the little stint, *Calidris minuta,* thereby assessing its potential as a bioindicator of environmental AMR contamination. To achieve this, we compared the resistance profiles of Enterobacterales and Gram-negative ESKAPE pathogens isolated from both *C. minuta* and their surrounding aquatic foraging sites characterized by differing levels of anthropogenic pressure.

## Materials and methods

### Study design and study area

This was a two-year cross-sectional study conducted in two distinct cohorts. The first cohort from *C. minuta* and water samples was collected in October 2021, coinciding with the arrival of *C. minuta* at the lakes from the Arctic. The second cohort of sample collection was in April 2022, during their departure from the lakes back to the Arctic. The study was conducted along the shores of Lake Magadi (02° 05′ 46″ S 36° 15′ 32″ E) and Lake Bogoria (02° 21′ 18″ N 36° 4′ 0″ E) and other surrounding aquatic ecosystems. Both lakes are saline-alkaline, are located in the arid Kenyan Rift Valley and are crucial wintering grounds for *C. minuta* (**Figs 1a** and **1b**) Lake Magadi, a closed basin in the southern Rift Valley near the Tanzanian border, is characterized by high concentrations of sodium chloride and carbonate compounds. Anthropogenic activity is substantial due to the commercial trona and soda ash mining industry. The local populations, majorly the pastoralists, Maasai, as well as tourists, utilize the hot (less than 45°C) spring pools for purported therapeutic benefits. In contrast, Lake Bogoria, a national reserve in the northern Rift Valley, is an open basin with comparatively lower concentrations of sodium chloride and carbonate. Human interactions with the lake are minimal, with the agro-pastoralists Tugens and animals (domestic and wildlife) accessing the freshwater springs by the lakes. Here, the hot springs, a major tourist attraction, are extremely hot (as high as 170°C), precluding human contact.

**Fig 1a.**
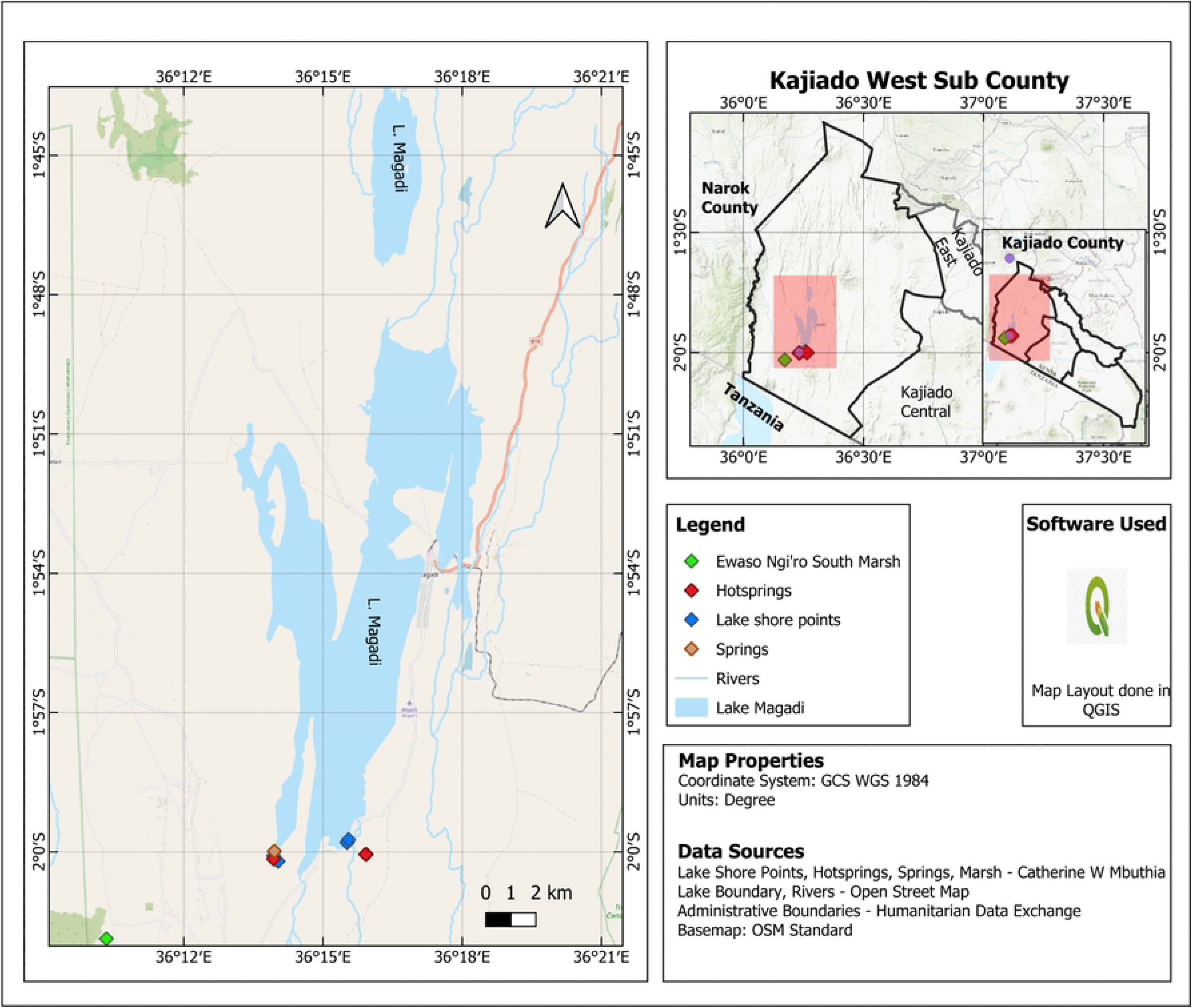
Map depicting sample collection sites for *C. minuta* and their aquatic foraging ecosystem sites in Lake Magadi.

**Fig 1b.**
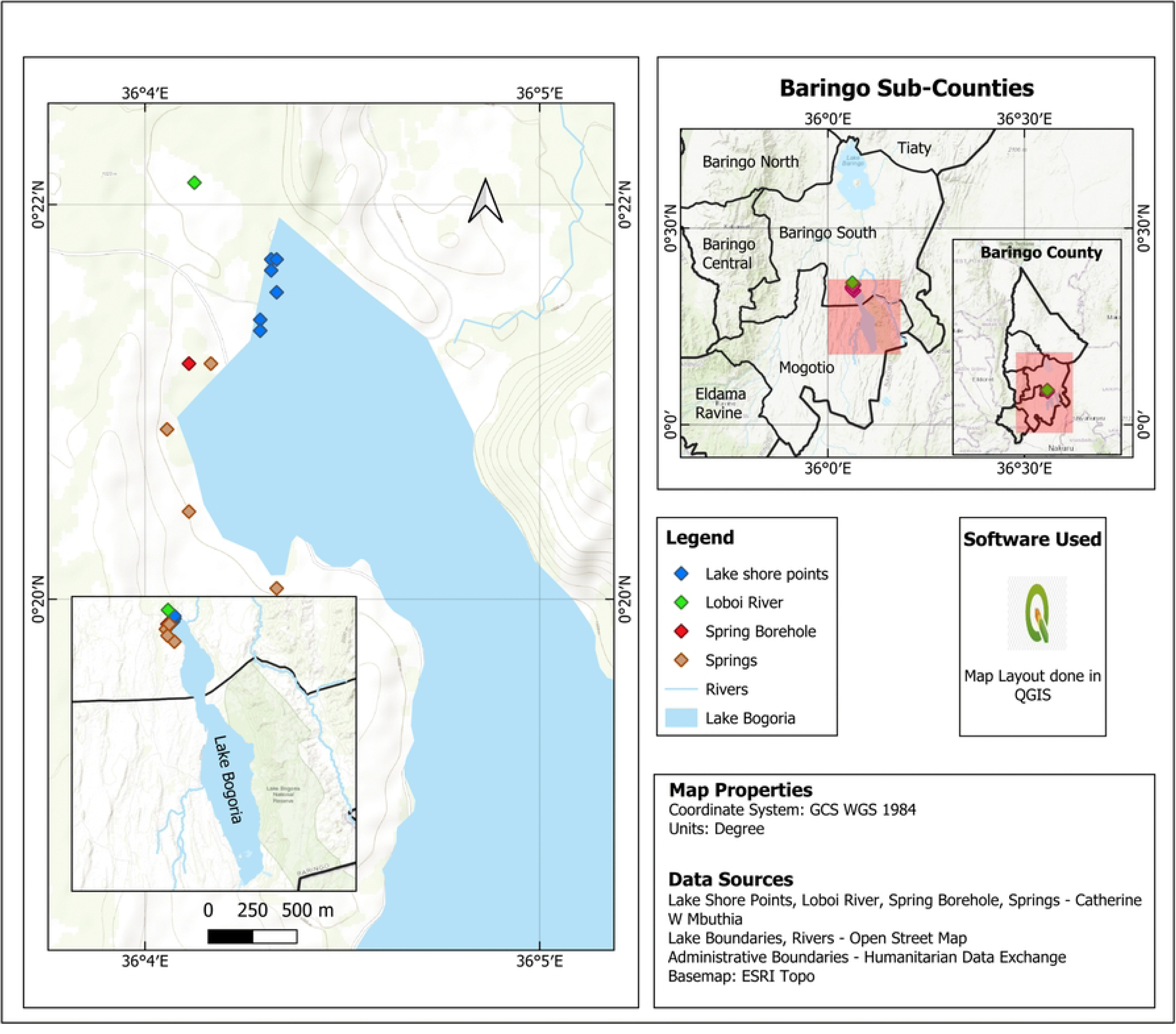
Map depicting sample collection sites for *C. minuta* and their aquatic foraging ecosystem sites in Lake Bogoria.

### Collection of samples

#### Sample collection from *C. minuta*

The birds were captured using mist nets placed in key areas along the lake shores, where humans and birds interact, such as hot springs, springs, lakes, and rivers. To prevent resampling, each bird was marked with a coded steel ring, and biometric information (age, health state, weight, moulting status) was recorded. A total of 184 non-invasive fresh fecal or cloacal samples were collected using sterile swabs from Lake Bogoria (cohort 1, n=45, and cohort 2, n=47) and Lake Magadi (cohort 1, n=36, and cohort 2, n=56). No anesthesia, euthanasia, or animal sacrifice was used on the birds. Samples were preserved in a Cary-Blair transport media (Oxoid, Basingstoke, UK) at -20° C during collection and processed within 72 h from collection.

#### Water sample collection

Water samples (n=48) were collected from various sites at Lake Bogoria (cohort 1, n=12, and cohort 2, n=13) and Lake Magadi (cohort 1, n=13, and cohort 2, n=12). The sampling locations included lakes, hot springs, rivers, and springs, which are part of the aquatic foraging environments of *C. minuta*. Approximately 50 mL of water was collected from each site and stored in sterile 50 mL Falcon tubes at -20° C during collection, and processed within 72 h from collection.

### Sample processing and laboratory procedures

#### Culture, isolation, and identification of Enterobacterales and ESKAPE pathogens

Following enrichment in buffered peptone water at 37°C for 18-24 hours, fecal or cloacal samples were streaked for isolation onto both MacConkey and Eosin Methylene Blue agars (Oxoid, Basingstoke, UK). For the water samples, aliquots of 50 mL were filtered using a Millipore vacuum filtration system apparatus, through filter papers of decreasing pore sizes (41μm, 20 μm, 10 μm, 1.2 μm, 0.8 μm, and finally 0.45 μm) (Thermo Fischer Scientific™ Nalgene™ Filter Membranes; Fischer Scientific, Hampton NH) to enable capture of all the bacteria present. All the filters were then aseptically placed into 50 mL MacConkey broth (Oxoid, Basingstoke, UK) in sterile Whirl packs. The inoculated broths were then thoroughly mixed and incubated at 37° C overnight. After incubation, each broth was streaked onto MacConkey (MAC) and Eosin Methylene Blue agar plates. After incubation at 37°C for a further 18-24 hours, suspected colonies were selected based on lactose fermentation patterns on MacConkey agar: early fermenters (pink colonies), late fermenters (less intense pink), and non-fermenters (colorless or pale colonies). Additional distinct morphological features were also used to target potential Enterobacterales and ESKAPE pathogens. A single pure colony from each suspect isolate was definitively identified using a Matrix-Assisted Laser Desorption/Ionization Time of Flight Mass Spectrometry (MALDI-TOF MS) system (BRUKER, Bremen, Germany) in accordance with the manufacturer’s protocol. All confirmed isolates were then preserved in tryptic soy broth with 15% glycerol and stored at - 80°C for subsequent analysis.

#### Phenotypic antimicrobial susceptibility testing

Antimicrobial susceptibility testing was conducted using the Kirby-Bauer disk diffusion method, following the Clinical and Laboratory Standards Institute guidelines (28). Previously identified isolates were recovered from frozen stock, sub-cultured on MacConkey agar, and then grown on Mueller-Hinton agar (Oxoid, Basingstoke, UK). A bacterial suspension adjusted to a 0.5 McFarland standard in sterile saline was uniformly lawned onto Mueller-Hinton agar plates.

A panel of 12 antimicrobial disks (Oxoid, Basingstoke, UK) from eight drug classes relevant to human and veterinary medicine and recommended for Enterobacteriaceae and ESKAPE pathogens, was applied. They included; ampicillin (AMP, 10 μg), amoxicillin-clavulanic acid (AMC, 30 (20:10) μg), cefepime (FEP, 30 μg), cefotaxime (CTX, 30 μg), ceftriaxone (CRO, 30 μg), cefuroxime (CXM, 30 μg), meropenem (MEM, 10 μg), ciprofloxacin (CIP, 5 μg), tetracycline (TE, 30 μg), chloramphenicol (C, 30 μg), gentamicin (CN, 10 μg), trimethoprim sulfamethoxazole (SXT, 25 (1.25: 23.75). Following incubation at 37°C for 18-24 hours, zones of inhibition were measured and interpreted as sensitive or resistant according to CLSI 2024 breakpoints.

Extended-spectrum beta-lactamase (ESBL) production was screened for via a double-disk synergy test (DDST) (28). Isolates displaying characteristic synergy zones between amoxicillin-clavulanate (AMC) and a third-generation cephalosporin were classified as ESBL producers. Multi-drug resistance (MDR) was defined as non-susceptibility to at least one agent in three or more antimicrobial categories (29). Quality control for the procedure was ensured using the *E. coli* ATCC 25922 reference strain.

### Statistical analyses

We employed a comprehensive comparative analysis to meticulously compare the occurrence of resistant bacterial isolates across four primary qualitative variables. The occurrence of resistant isolates was compared between (i) Sample Source: *C. minuta* and water samples (collected at the birds’ trapping sites), (ii) Study Areas: High and low anthropogenic intensification (Lakes Magadi and Bogoria), respectively and (iii) Cohorts: Cohort 1 (coinciding with the birds’ arrival from the Arctic) and Cohort 2 (birds’ departure from the Rift Valley lakes). Residuals of resistance rates from one-way ANOVA by the qualitative variables (sample source, antimicrobials, study area, and cohorts) were subjected to the Shapiro-Wilk test to assess their normality. This yielded a p-value of 0.0046, indicating that resistance rates were non-normally distributed, and therefore non-parametric tests would be more appropriate. Consequently, a tie-corrected Kruskal-Wallis test was employed to test for differences in medians of resistance rates across the four qualitative variables separately.

### Ethical considerations

The study obtained scientific and ethical clearances from the National Commission for Science Technology, and Innovation (NACOSTI), identification number 333849, and License No: NACOSTI/P/21/13520, and the Directorate of Postgraduate Studies, Research, Technology Transfer and Consultancy (DPRTC) at the Sokoine University of Agriculture. Site access and wildlife permits for collecting *C. minuta* samples were granted by the department of gender, culture, tourism, and wildlife of Kajiado county government and department of environment, natural resources, tourism, and wildlife of Baringo county government. Water sample collection approvals were provided by the department of water, environment, and natural resources and the department of water, irrigation, environment, natural resources, climate change, and mining of Kajiado and Baringo County governments, respectively.

## Results

### Identification and characterization of bacterial isolates in *C. minuta* and water samples

A total of 294 bacterial isolates encompassing 16 genera and 33 distinct species were recovered. The *C. minuta* samples yielded 233 isolates (76 from Bogoria, 157 from Magadi), while the water samples provided 61 isolates (34 from Bogoria, 27 from Magadi). A notable similarity was observed in the diversity of Enterobacterales and Gram-negative ESKAPE species between the bird and water samples at both lakes (**Figs 2a** and **2b**). The most frequently isolated species were Enterobacter species (31.0%) and *Escherichia coli* (17.3%).

**Fig 2a.**
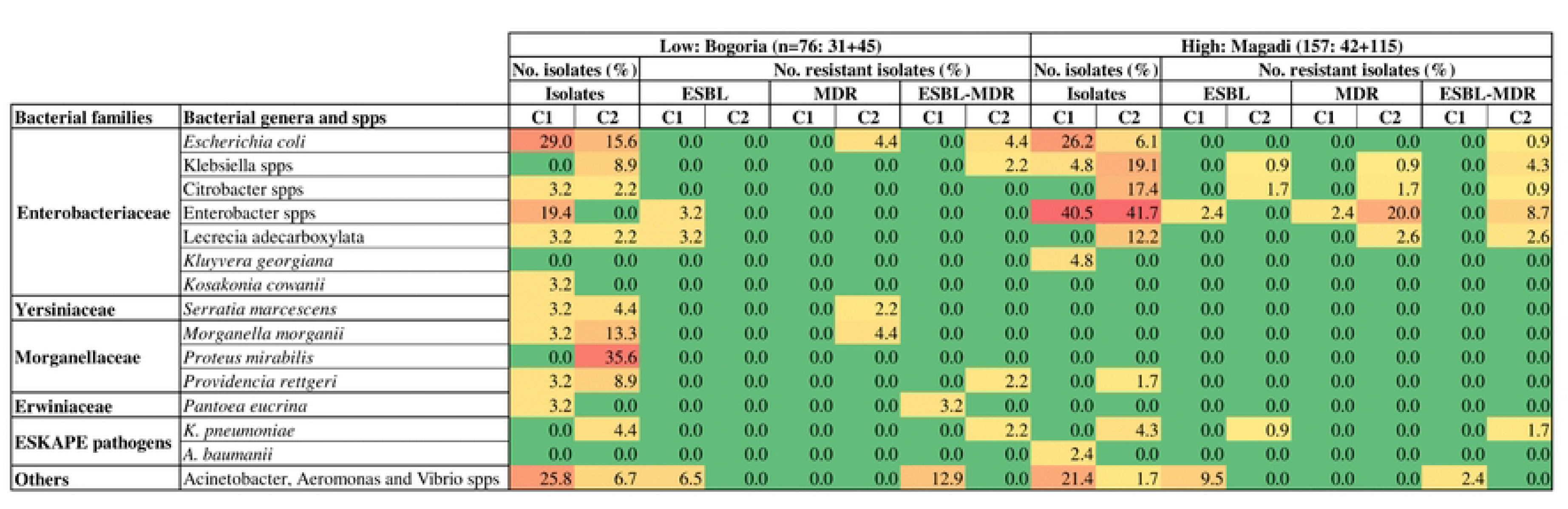
Heat map depicting the frequency (%) of bacterial isolates and their resistances in *C. minuta* from areas of low (Lake Bogoria) and high (Lake Magadi) anthropogenic intensification. Different color bars indicate different resistance frequencies. Red: High, Yellow: Moderate, Green: Low

**Fig 2b.**
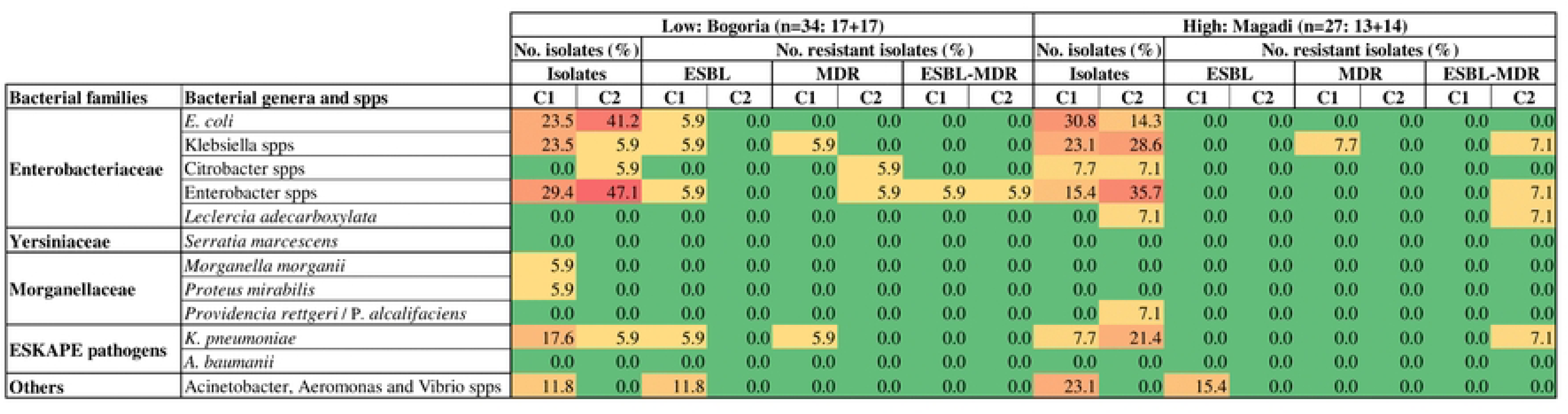
Heat map depicting the frequency (%) of bacterial isolates and their resistances in water from areas of low (Lake Bogoria) and high (Lake Magadi) anthropogenic intensification. Different color bars indicate different resistance frequencies. Red: High, Yellow: Moderate, Green: Low

Analysis also identified bacteria beyond the primary Enterobacterales and ESKAPE targets, classified as ‘Others’ (**Figs 2a** and **2b**). These isolates, predominantly recovered from *C. minuta*, belonged to the orders Pseudomonadales (*Acinetobacter soli*, *A. hemolyticus*, and *A. balylyi*), Aeromonadales (*Aeromonas veronii* and *A. hydrophila*), and Vibrionales (*Vibrio metschnikovii* and *V. albensis*).

### Frequency and distribution of multi-drug-resistant and extended-spectrum beta-lactamase-producing bacterial isolates

Of the 294 bacterial isolates analyzed, the frequency of multi-drug resistant (MDR), extended-spectrum beta-lactamases (ESBLs) and ESBL-MDR phenotypes varied among the species and across the two sampling sites, sources and cohorts as illustrated in **Figs 2a** and **2b**. Enterobacter species (8.5%) displayed the highest frequency of MDR, followed by Klebsiella species (1%). The highest rates of ESBLs-production were in Acinetobacter species (3.4%) and Enterobacter species (1.0%). Furthermore, the ESBL-MDR phenotypes were more prevalent in Enterobacter species (4.4%) followed by Klebsiella species (2.4%) and *Escherichia coli* (1.0%). A notable finding was the frequency of antimicrobial resistance among ‘Others’ isolates-the Acinetobacter species. From Lake Bogoria, *C. minuta* yielded five Acinetobacter strains (5/76, 6.6%) with an ESBL-MDR phenotype and one (1/76, 1.3%) as an ESBL-producer. Water samples from this lake contained two additional ESBL-producing *A. soli* strains (2/34, 5.9%). At Lake Magadi, *C. minuta* carried four *A. balylyi* strains (4/157, 2.5%) with ESBL phenotypes and one with an ESBL-MDR phenotype, while water samples yielded two further ESBL-producing *A. balylyi* strains (2/27. 7.4%).

### Antimicrobial resistance profiles

Resistance profiles varied considerably between sample groups and cohorts. The most widespread resistances were to ampicillin (50.0% of isolates), amoxicillin-clavulanic acid (36.4%), tetracycline (32.7%), trimethoprim-sulfamethoxazole (24.1%), and cefotaxime (18.4%). In contrast, susceptibility remained high for meropenem (1.0% resistance) and cefepime (3.4% resistance). A complete summary of resistance frequencies for all tested antimicrobials, stratified by cohort, site, and sample group, is provided in **Table 1**.

**Table 1.** Antimicrobial resistance frequencies (%) of Enterobacterales and ESKAPE pathogens isolated from *C. minuta* and water samples in areas of low (Lake Bogoria) and high (Lake Magadi) anthropogenic intensification.

| Antimicrobial | *C.minuta* |  | Water |  | Total |
| --- | --- | --- | --- | --- | --- |
|  | L. Bogoria | L. Magadi | L. Bogoria | L. Magadi |  |

|  | C1<br>(n=31) | C2<br>(n=45) | C1<br>(n=42) | C2<br>(n=115) | C1<br>(n=17) | C2<br>(n=17) | C1<br>(n=13) | C2<br>(n=14) | n=294 |
| --- | --- | --- | --- | --- | --- | --- | --- | --- | --- |
| AMP | 8(25.8) | 15(33.3) | 13(31.0) | 86(74.8) | 5(29.4) | 9(52.9) | 5(38.5) | 6(42.9) | 147(50) |
| AMC | 6(19.4) | 7(15.6) | 7(16.7) | 70(60.1) | 6(35.3) | 5(29.4) | 0(0.0) | 6(42.9) | 107(36.4) |
| FEP | 4(12.9) | 3(6.7) | 0(0.0) | 2(1.8) | 0(0.0) | 0(0.0) | 0(0.0) | 1(7.1) | 10(3.4) |
| CTX | 9(29.0) | 4(8.9) | 6(14.3) | 23(20) | 6(35.3) | 1(5.9) | 2(15.4) | 3(21.4) | 54(18.4) |
| CRO | 5(16.1) | 3(6.7) | 4(9.5) | 2(1.8) | 2(11.8) | 0(0.0) | 0(0.0) | 1(7.1) | 17(5.8) |
| CXM | 4(12.9) | 5(11.1) | 0(0.0) | 3(2.6) | 1(5.9) | 1(5.9) | 0(0.0) | 0(0.0) | 14(4.8) |
| MEM | 2(6.5) | 0(0.0) | 0(0.0) | 0(0.0) | 0(0.0) | 0(0.0) | 0(0.0) | 1(7.1) | 3(1.0) |
| CIP | 6(19.4) | 0(0.0) | 2(4.8) | 8(7.0) | 0(0.0) | 0(0.0) | 0(0.0) | 3(21.4) | 19(6.5) |
| TE | 2(6.5) | 23(51.1) | 4(9.5) | 54(47) | 2(11.8) | 2(11.8) | 5(38.5) | 4(28.6) | 96(32.7) |
| C | 2(6.5) | 5(11.1) | 0(0.0) | 7(6.1) | 0(0.0) | 0(0.0) | 0(0.0) | 0(0.0) | 14(4.8) |
| CN | 1(3.2) | 3(6.7) | 1(2.4) | 4(3.5) | 1(5.9) | 1(5.9) | 0(0.0) | 2(14.3) | 13(4.4) |
| SXT | 1(3.2) | 6(13.3) | 3(7.1) | 53(46.1) | 0(0.0) | 0(0.0) | 4(30.8) | 4(28.6) | 71(24.1) |

### Statistical analyses

Through a tie-corrected Kruskal-Wallis Test comparison of medians, there were more resistant strains from *C. minuta* than from the water isolates. In addition, isolates from Lake Magadi showed higher resistance frequencies than those from Lake Bogoria. Results from the two cohorts showed that isolates from cohort 2 recorded higher rates of resistance than those from cohort 1. It was also observed that ampicillin, amoxicillin-clavulanic acid, and tetracycline showed the highest resistance in comparison to other antimicrobials. On the contrary, meropenem and cefepime recorded the best susceptibility rates (**Table 2**).

**Table 2.** Resistance frequencies of bacterial isolates: Descriptive statistics and comparative analysis using a Tie-corrected Kruskal-Wallis Test.

| Variable | N | Median (IQR) | Min-Max | K-Wallis Test* |
| --- | --- | --- | --- | --- |
| **Source** | | | | $\chi^2$ (1) = 2.105<br>P-value = 0.1468 |
| *C.minuta* | 48 | 8.00(14.85) | 0.00- 74.80 |  |
| Water | 48 | 5.90(25.00) | 0.00 - 52.90 |  |
| **Antimicrobials** | | | | $\chi^2$ (11) = 50.653<br>P-value = 0.0001 |
| AMC | 8 | 24.40(22.95) | 0.00 - 60.10 |  |
| AMP | 8 | 35.90(17.70) | 25.80 - 74.80 |  |
| C | 8 | 0.00(6.3) | 0.00 - 11.10 |  |
| CIP | 8 | 2.40(13.20) | 0.00 - 21.40 |  |
| CN | 8 | 4.70(3.50) | 0.00 - 14.30 |  |
| CRO | 8 | 6.90(9.75) | 0.00 - 16.10 |  |
| CTX | 8 | 17.70(13.60) | 5.90 - 35.30 |  |
| CXM | 8 | 4.25(8.50) | 0.00 - 12.90 |  |
| FEP | 8 | 0.90(6.90) | 0.00 - 12.90 |  |
| MEM | 8 | 0.00(3.25) | 0.00 - 7.10 |  |
| SXT | 8 | 10.20(28.10) | 0.00 - 46.10 |  |
| TE | 8 | 20.20(32.10) | 6.50 - 51.10 |  |
| **Study area** | | | | $\chi^2 (1) = 0.045$<br>P-value = 0.8325 |
| L. Bogoria | 48 | 6.70(15.85) | 0.00 - 52.90 |  |
| L. Magadi | 48 | 7.10(25.00) | 0.00 - 74.80 |  |
| **Cohort** | | | | $\chi^2 (1) = 1.573$<br>P-value = 0.2098 |
| C1 | 48 | 6.50(16.40) | 0.00 - 38.50 |  |
| C2 | 48 | 7.10(23.20) | 0.00 - 74.80 |  |
| **Total** | 96 | 6.85(19.70) | 0.00 - 74.80 |  |

\* Tie-corrected Kruskal–Wallis test between resistance frequencies and each variable.

**Table 2** summarizes the distribution of resistance frequencies across sample source, study areas, antimicrobials, and cohorts using medians and interquartile ranges (IQR). The Kruskal– Wallis test indicated that there were no significant differences in median rates: between *C. minuta* and water isolates (χ² (1) = 2.11, p = 0.147); Lake Bogoria and Lake Magadi (χ² (1) = 0.045, p = 0.833); and between cohorts 1and 2 (χ² (1) = 1.57, p = 0.210). However, there were significant differences across the 12 antimicrobials (χ² (11) = 50.65, p < 0.0001). Thus, antimicrobials were the only factor strongly associated with variation in resistance frequencies.

## Discussion

Our findings align with established research linking anthropogenic activities to the prevalence of antimicrobial resistance (AMR) in wild birds. Numerous studies have documented this connection in contexts such as intensive agriculture (30) (13), landfills and dump sites (19) (31), urban areas (32), industries (33), wastewater treatment systems (34) (35). The higher bacterial resistance observed in *C. minuta* foraging at the heavily impacted Lake Magadi, compared to the less disturbed Lake Bogoria, corroborates this pattern.

Aquatic environments are considered primary transmission routes for AMR bacteria to wild birds, a phenomenon demonstrated by (35). This risk is amplified for *C. minuta* at Lake Magadi, where a bird’s specific foraging ecology exposes it to multiple contamination sources. These include hot springs frequented by humans, the Ewaso Ng’iro South River shared with livestock, and commercial salt pans producing industrial effluent. This multi-faceted anthropogenic pressure is reflected in the isolation of similar, often human and livestock-associated resistant strains from both the birds and their aquatic foraging ecosystems.They included isolates associated with opportunistic human pathogens such as *Lecrecia adecarboxylata* (36) *, Kluyvera georgiana* (37), and multi-drug resistant *Acinetobacter spps* (38). In contrast, the lower AMR frequencies at Lake Bogoria correspond with its limited human contact, dominated by wildlife (crocodiles and hippopotamuses). However, the detection of resistant, human-originated *Acinetobacter* pathogens even in this less-impacted site indicates that resistant strains are now ubiquitous in the environment. This ubiquity suggests that AMR is ‘embedded’ within natural ecosystems, enabling migratory birds to acquire human-associated resistance far from direct sources of contamination. This presents a significant challenge for One Health surveillance, complicating efforts to trace AMR exchanges and track the global spread of emerging resistance genes (39) (40).

The diversity of resistant Enterobacterales and ESKAPE pathogens isolated from *C. minuta* is consistent with reports from other wild avian populations (25), (27) (26), supporting the role of wild birds in the transboundary transmission of AMR. The slightly higher resistance frequencies in the cohort sampled prior to departure from the lakes (Cohort 2) compared to the arriving Arctic cohort (Cohort 1) suggest accumulation of resistant strains during their stay, though the difference was not statistically significant. This implies exposure to environmental pollution across their entire annual migratory cycle.

Of particular concern is the high prevalence of resistance in *C. minuta* to ampicillin, amoxicillin-clavulanic acid, tetracycline, and trimethoprim-sulfamethoxazole-antimicrobials heavily used in human and veterinary medicine. The significant resistance to the World Health Organization (WHO) critically important antimicrobials (CIA) List for human medicine (WHO CIA), such as cefotaxime and ciprofloxacin, and the notable presence of ESBL-producing strains, is alarming. The location of these resistance genes on mobile genetic elements (MGEs) facilitates their rapid horizontal transfer across bacterial species and ecosystems (15), accelerating the proliferation of ESBLs within the One Health framework (41) (42) (43). The concurrent isolation of MDR strains further narrows the spectrum of effective treatments. While resistance to last-resort drugs like cefepime and meropenem remains low, the ongoing global misuse of antimicrobials risks fostering resistance to these final-line agents in wild bird populations, ultimately depleting our therapeutic arsenal.

## Conclusion

These findings highlight the impact of anthropogenic activities and the crucial role of migratory *C. minuta* as reservoirs and potential accelerators in the global dissemination of AMR. By disseminating these superbugs across continents during their annual migrations, these birds amplify AMR spread. Since wild birds are not directly subjected to antimicrobials therapies, effective management of AMR in these populations requires a different strategy: proactive surveillance and interventions through a holistic One-Health framework. This necessity is fundamentally aligned with the “Zero Pollution Vision for 2050, which recognizes environmental pollution as a catalyst for AMR (European Commission, 2021). Implementing environmental stewardship measures to safeguard avian populations from AMR will simultaneously achieve broader One Health objectives and contribute to a healthier global ecosystem.

## Acknowledgements

We acknowledge and express gratitude to the Center for Microbiology at the Kenya Medical Research Institute (CMR-KEMRI), Sokoine University of Agriculture, SACIDS Foundation for One Health, the International Livestock Research Institute and Karatina University for guidance, infrastructure, and laboratory supplies. We also specifically thank Gerishom Angote, Benard Ochiel, and Hulder Otieno of the Centre for Microbiology at KEMRI for their immense laboratory input.

## References

1. Ahmed SK, Hussein S, Qurbani K, Ibrahim RH, Fareeq A, Mahmood KA, et al. Antimicrobial resistance: Impacts, challenges, and future prospects. J Med Surg Public Health. 2024 Apr; 2:100081.

2. Collignon PJ, McEwen SA. One Health—Its Importance in Helping to Better Control Antimicrobial Resistance. Trop Med Infect Dis. 2019 Mar;4(1):22.

3. Thakur S, Gray GC. The Mandate for a Global “One Health” Approach to Antimicrobial Resistance Surveillance. Am J Trop Med Hyg. 2019 Feb 1;100(2):227–8.

4. D’Costa VM, King CE, Kalan L, Morar M, Sung WWL, Schwarz C, et al. Antibiotic resistance is ancient. Nature. 2011 Sep;477(7365):457–61.

5. Allel K, Day L, Hamilton A, Lin L, Furuya-Kanamori L, Moore CE, et al. Global antimicrobial-resistance drivers: an ecological country-level study at the human–animal interface. Lancet Planet Health. 2023 Apr;7(4):e291–303.

6. Manyi-Loh C, Mamphweli S, Meyer E, Okoh A. Antibiotic Use in Agriculture and Its Consequential Resistance in Environmental Sources: Potential Public Health Implications. Molecules. 2018 Apr;23(4):795.

7. Roth N, Käsbohrer A, Mayrhofer S, Zitz U, Hofacre C, Domig KJ. The application of antibiotics in broiler production and the resulting antibiotic resistance in Escherichia coli: A global overview. Poult Sci. 2019 Apr;98(4):1791–804.

8. Blanco G, Bautista LM. Avian Scavengers as Bioindicators of Antibiotic Resistance Due to Livestock Farming Intensification. Int J Environ Res Public Health. 2020 Jan;17(10):3620.

9. Oyelayo EA, Taiwo TJ, Oyelude SO, Alao JO. The global impact of industrialisation and climate change on antimicrobial resistance: assessing the role of Eco-AMR Zones. Environ Monit Assess. 2025 May 5;197(6):625.

10. Padma K. Overuse and Misuse of Antibiotics. J Biomed Pharm Res [Internet]. 2022 Mar 10 [cited 2025 May 10];11(1). Available from: http://jbpr.in/index.php/jbpr/article/view/899

11. Caneschi A, Bardhi A, Barbarossa A, Zaghini A. The Use of Antibiotics and Antimicrobial Resistance in Veterinary Medicine, a Complex Phenomenon: A Narrative Review. Antibiotics. 2023 Mar 1;12(3):487.

12. Hasan B, Melhus Å, Sandegren L, Alam M, Olsen B. The Gull ( *Chroicocephalus brunnicephalus* ) as an Environmental Bioindicator and Reservoir for Antibiotic Resistance on the Coastlines of the Bay of Bengal. Microb Drug Resist. 2014 Oct;20(5):466–71.

13. Elsohaby I, Samy A, Elmoslemany A, Alorabi M, Alkafafy M, Aldoweriej A, et al. Migratory Wild Birds as a Potential Disseminator of Antimicrobial-Resistant Bacteria around Al-Asfar Lake, Eastern Saudi Arabia. Antibiotics. 2021 Mar 5;10(3):260.

14. Guardia T, Varialle L, Mnichino A, Balestrieri R, Mastronardi D, Russo T, et al. Wild birds and the ecology of antimicrobial resistance: an approach to monitoring. 2024 Apr 24;e22588.

15. Báez J, Hernández-García M, Guamparito C, Díaz S, Olave A, Guerrero K, et al. Molecular Characterization and Genetic Diversity of ESBL-Producing Escherichia coli Colonizing the Migratory Franklin’s Gulls (Leucophaeus pipixcan) in Antofagasta, North of Chile. Microb Drug Resist. 2015 Feb;21(1):111–6.

16. Bouaziz A, Loucif L, Ayachi A, Guehaz K, Bendjama E, Rolain JM. Migratory White Stork (Ciconia ciconia): A Potential Vector of the OXA-48-Producing Escherichia coli ST38 Clone in Algeria. Microb Drug Resist. 2018 May;24(4):461–8.

17. Ahmed ZS, Elshafiee EA, Khalefa HS, Kadry M, Hamza DA. Evidence of colistin resistance genes (mcr-1 and mcr-2) in wild birds and its public health implication in Egypt. Antimicrob Resist Infect Control. 2019;8:197.

18. Yuan Y, Liang B, Bo-wen J, Ling-wei Z, Tie-cheng W, Yuan-guo L, et al. Migratory wild birds carrying multidrug-resistant Escherichia coli as potential transmitters of antimicrobial resistance in China. 2021 Dec;e0261444.

19. Bonnedahl J, Hernandez J, Stedt J, Waldenström J, Olsen B, Drobni M. Extended-Spectrum β-Lactamases in *Escherichia coli and Klebsiella pneumoniae* in Gulls, Alaska, USA. Emerg Infect Dis [Internet]. 2014 May [cited 2024 Dec 8];20(5). Available from: http://wwwnc.cdc.gov/eid/article/20/5/13-0325_article.htm

20. Tarabai H, Krejci S, Karyakin I, Bitar I, Literak I, Dolejska M. Clinically relevant antibiotic resistance in Escherichia coli from black kites in southwestern Siberia: a genetic and phenotypic investigation. mSphere. 2023 Jun 13;8(4):e00099–23.

21. Waldenström J, Toor M van, Lindström Å. Long-term trends in abundance, phenology, and morphometrics of Little Stint Calidris minuta during autumn migration in southern Sweden, 1946–2020. Ornis Svec. 2023 Jun 1;33:30–48.

22. Yosef R, Gołdyn B, Zduniak P. Predation of migratory Little Stint ( Calidris minuta ) by Barbary Falcon ( Falco pelegrinoides ) is dependent on body mass and duration of stopover time. 2011;257–61.

23. Pearson DJ. The status, migrations and seasonality of the Little Stint in Kenya. Ringing Migr. 1987 Oct;8(2):91–108.

24. Foti M, Mascetti A, Fisichella V, Fulco E, Orlandella BM. Antibiotic resistance assessment in bacteria isolated in migratory Passeriformes transiting through the Metaponto territory. Avian Res. 2017;1–11.

25. Machado D, Lopes E, Albuquerque A, Horn R, Bezerra W, Siqueira R, et al. Isolation and Antimicrobial Resistance Profiles of Enterobacteria from Nestling Grey-Breasted Parakeets (Pyrrhura Griseipectus). Rev Bras Ciênc Avícola. 2018 Mar;20(1):103–10.

26. Russo TP, Minichino A, Gargiulo A, Varriale L, Borrelli L, Pace A, et al. Prevalence and Phenotypic Antimicrobial Resistance among ESKAPE Bacteria and Enterobacterales Strains in Wild Birds. Antibiotics. 2022 Dec 15;11(12):1825.

27. Darwich L, Vidal A, Seminati C, Albamonte A, Casado A, López F, et al. High prevalence and diversity of Extended-Spectrum β-Lactamase and emergence of Carbapenemase producing Enterobacteriaceae spp in wildlife in Catalonia [Internet]. bioRxiv; 2019 [cited 2022 Dec 23]. p. 510123. Available from: https://www.biorxiv.org/content/10.1101/510123v1

28. CLSI M100, 2024. Clinical and Laboratory Standards Institute (CLSI). Performance Standards for Antimicrobial Susceptibility Testing, 34th ed.; CLSI: Wayne, PA, USA; 2024.

29. Magiorakos AP, Srinivasan A, Carey RB, Carmeli Y, Falagas ME, Giske CG, et al. Multidrug-resistant, extensively drug-resistant and pandrug-resistant bacteria: An international expert proposal for interim standard definitions for acquired resistance. Clin Microbiol Infect. 2012;18(3):268–81.

30. Chandler JC, Anders JE, Blouin NA, Carlson JC, LeJeune JT, Goodridge LD, et al. The Role of European Starlings (Sturnus vulgaris) in the Dissemination of Multidrug-Resistant Escherichia coli among Concentrated Animal Feeding Operations. Sci Rep. 2020 May 15;10(1):8093.

31. Merkeviciene L, Klimiene I, Siugzdiniene R, Virgailis M, Mockeliunas R, Ruzauskas M. Prevalence and molecular characteristics of multi-resistant *Escherichia coli* in wild birds. Acta Vet Brno. 2018 Apr 12;87(1):9–17.

32. Ben Yahia H, Ben Sallem R, Tayh G, Klibi N, Ben Amor I, Gharsa H, et al. Detection of CTX-M-15 harboring Escherichia coli isolated from wild birds in Tunisia. BMC Microbiol. 2018 Apr 2;18(1):26.

33. Stedt J, Bonnedahl J, Hernandez J, McMahon BJ, Hasan B, Olsen B, et al. Antibiotic resistance patterns in Escherichia coli from gulls in nine European countries. Infect Ecol Epidemiol. 2014 Jan 1;4(1):21565.

34. Hasan B, Laurell K, Rakib MM, Ahlstedt E, Hernandez J, Caceres M, et al. Fecal Carriage of Extended-Spectrum β-Lactamases in Healthy Humans, Poultry, and Wild Birds in León, Nicaragua—A Shared Pool of blaCTX-M Genes and Possible Interspecies Clonal Spread of Extended-Spectrum β-Lactamases-Producing Escherichia coli. Microb Drug Resist. 2016 Dec;22(8):682–7.

35. Wu J, Huang Y, Rao D, Zhang Y, Yang K. Evidence for Environmental Dissemination of Antibiotic Resistance Mediated by Wild Birds. Front Microbiol [Internet]. 2018 [cited 2023 Jul 27];9(745). Available from: https://www.frontiersin.org/articles/10.3389/fmicb.2018.00745

36. Zayet S, Lang S, Garnier P, Pierron A, Plantin J, Toko L, et al. Leclercia adecarboxylata as Emerging Pathogen in Human Infections: Clinical Features and Antimicrobial Susceptibility Testing. Pathogens. 2021 Oct 28;10(11):1399.

37. Torre D, Crespi E, Bernasconi M, Rapazzini P. Urinary tract infection caused by Kluyvera ascorbata in an immunocompromised patient: Case report and review. Scand J Infect Dis. 2005 May;37(5):375–8.

38. Chen TL, Siu LK, Lee YT, Chen CP, Huang LY, Wu RCC, et al. *Acinetobacter baylyi* as a Pathogen for Opportunistic Infection. J Clin Microbiol. 2008 Sep;46(9):2938–44.

39. Vittecoq M, Godreuil S, Prugnolle F, Durand P, Brazier L, Renaud N, et al. Antimicrobial resistance in wildlife. 2016 [cited 2019 May 31]; Available from: https://access.webofknowledge.com/

40. Laborda P, Sanz-García F, Ochoa-Sánchez LE, Gil-Gil T, Hernando-Amado S, Martínez JL. Wildlife and Antibiotic Resistance. Front Cell Infect Microbiol. 2022 May 11;12:873989.

41. Fahim KM, Ismael E, Khalefa HS, Farag HS, Hamza DA. Isolation and characterization of E. coli strains causing intramammary infections from dairy animals and wild birds. Int J Vet Sci Med. 2019 Jan 2;7(1):61–70.

42. Ngaiganam EP, Pagnier I, Chaalal W, Leangapichart T, Chabou S, Rolain JM, et al. Investigation of urban birds as source of β-lactamase-producing Gram-negative bacteria in Marseille city, France. Acta Vet Scand. 2019 Oct 31;61(1):51.

43. Belmahdi M, Chenouf NS, Ait Belkacem A, Martinez-Alvarez S, Pino-Hurtado MS, Benkhechiba Z, et al. Extended Spectrum β-Lactamase-Producing Escherichia coli from Poultry and Wild Birds (Sparrow) in Djelfa (Algeria), with Frequent Detection of CTX-M-14 in Sparrow. Antibiotics. 2022 Dec;11(12):1814.

44. Commission E. Wilkki, C.M., and Reeve, N. (2021). Communication from the Commission to the European Parliament, the Council, the European Economic and Social Committee and the Committee of the Regions on European Missions European Commission Directorate-General for Research and Innovation Directorate G—Common Policy Centre. European Commission: Brussels, Belgium. 2021.

